# Developmental maturation of the precuneus as a functional core of the default-mode network

**DOI:** 10.1101/419028

**Authors:** Rosa Li, Amanda V. Utevsky, Scott A. Huettel, Barbara R. Braams, Sabine Peters, Eveline A. Crone, Anna C.K. van Duijvenvoorde

## Abstract

Efforts to map the functional architecture of the developing human brain have shown that connectivity between and within functional neural networks changes from childhood to adulthood. While prior work has established that the adult precuneus distinctively modifies its connectivity during task versus rest states (Utevsky, Smith, & Huettel, 2014), it remains unknown how these connectivity patterns emerge over development. Here, we use functional magnetic resonance imaging (fMRI) data collected at two longitudinal timepoints from over 250 participants between the ages of 8 and 26 engaging in two cognitive tasks and a resting-state scan. By applying independent component analysis (ICA) to both task and rest data, we identified three canonical networks of interest – the rest-based default mode network (DMN) and the task-based left and right frontoparietal networks (LFPN, RFPN) – which we explored for developmental changes using dual-regression analyses. We found systematic state-dependent functional connectivity in the precuneus, such that engaging in a task (compared to rest) resulted in greater precuneus-LFPN and precuneus-RFPN connectivity, whereas being at rest (compared to task) resulted in greater precuneus-DMN connectivity. These cross-sectional results replicated across both tasks and at both developmental timepoints. Finally, we used longitudinal mixed models to show that the degree to which precuneus distinguishes between task and rest states increases with age, due to age-related increasing segregation between precuneus and LFPN at rest. Our results highlight the distinct role of the precuneus in tracking processing state, in a manner that is both present throughout and strengthened across development.

## Introduction

The human brain exhibits distinct patterns of functional connectivity between disparate brain regions, even when at rest. These patterns are so reliably evoked across studies and participants that they can be described as a set of canonical neural networks reflecting the intrinsic functional organization of the human brain (Smith et al., 2009; van den Heuvel & Hulshoff Pol, 2010). The default-mode network (DMN), comprised of the precuneus, posterior cingulate cortex, medial prefrontal cortex, and bilateral temporoparietal junction, is the network most readily associated with rest states, as its activity increases during rest and decreases during task engagement (Raichle et al., 2001; Shulman et al., 1997). Other networks, however, also show patterns of intrinsic connectivity during rest, including lateralized frontoparietal networks (FPNs) that are more generally associated with task-positive, goal-directed attention (Corbetta & Shulman, 2002; Vincent, Kahn, Snyder, Raichle, & Buckner, 2008) and found to be anti-correlated with the DMN (Fox et al., 2005).

Within the regions of the DMN, the precuneus stands out for its distinctive role. Several studies have shown that, despite being a component of the DMN, precuneus activation increases during tasks such as memory retrieval (Fletcher et al., 1995; Lundstrom, Ingvar, & Petersson, 2005; Maddock, Garrett, & Buonocore, 2001), reward monitoring (Hayden, Nair, McCoy, & Platt, 2008), and emotion processing (Maddock, Garrett, & Buonocore, 2003); see (Cavanna & Trimble, 2006) for review). Notably, Utevsky, Smith, & Huettel (2014) found precuneus to be the only neural region that both increased connectivity with DMN at rest compared to task *and* increased connectivity with left FPN (LFPN) at task compared to rest. While greater precuneus-DMN connectivity during rest may be expected as a result of increased within-network connectivity, the finding of greater precuneus-LFPN connectivity during task is more counterintuitive, as precuneus is not part of the LFPN. These results suggest that precuneus serves as a functional core of the DMN by altering its network connectivity to LFPN and DMN according to whether the brain is in a task or a rest state.

While precuneus connectivity has been studied using data from adults (Honey et al., 2009; Leech, Braga, & Sharp, 2012; Leech, Kamourieh, Beckmann, & Sharp, 2011; Utevsky et al., 2014), the role of the precuneus in mediating between task and rest states has not yet been investigated across development. Previous work using resting-state data has shown that within-network functional and structural connectivity increases with age for DMN and FPNs (Baum et al., 2017; Fair et al., 2007, 2008; Uddin, Supekar, Ryali, & Menon, 2011), and that the DMN and FPNs become increasingly segregated from each other with age (Sherman et al., 2014), consistent with the idea that within-network connections are strengthened and between-network connections are weakened across development (Fair et al., 2009). Therefore we expected that the precuneus, as a node of the DMN, would show developmental changes in connectivity with DMN and FPN, reflecting change in the degree to which this functional region mediates between task and rest states.

Here, we used network-based connectivity analyses to probe the role of the precuneus in a large accelerated longitudinal sample. Over 250 participants between the ages of 8 and 26 completed a resting state scan and two cognitive task scans at two longitudinal timepoints approximately two years apart. We used data from the first longitudinal timepoint to identify rest and task-based networks of interest, DMN and left and right FPN (RFPN). We then used dual-regression analyses (Filippini et al., 2009; R. Leech et al., 2012; Robert Leech et al., 2011; Nickerson, Smith, Öngür, & Beckmann, 2017; Smith et al., 2014) to examine task versus rest connectivity with our networks of interest at both longitudinal timepoints and for both tasks (see Figure 1 for a schematic analysis diagram). This design allowed us to self-replicate connectivity results across two longitudinal timepoints, and self-replicate developmental results across two different tasks. Our results extended previous findings in adults (Utevsky et al., 2014) to children and adolescents, while also testing the hypothesis that the role of the precuneus as a functional core of the DMN strengthens across human development.

**Figure 1:**
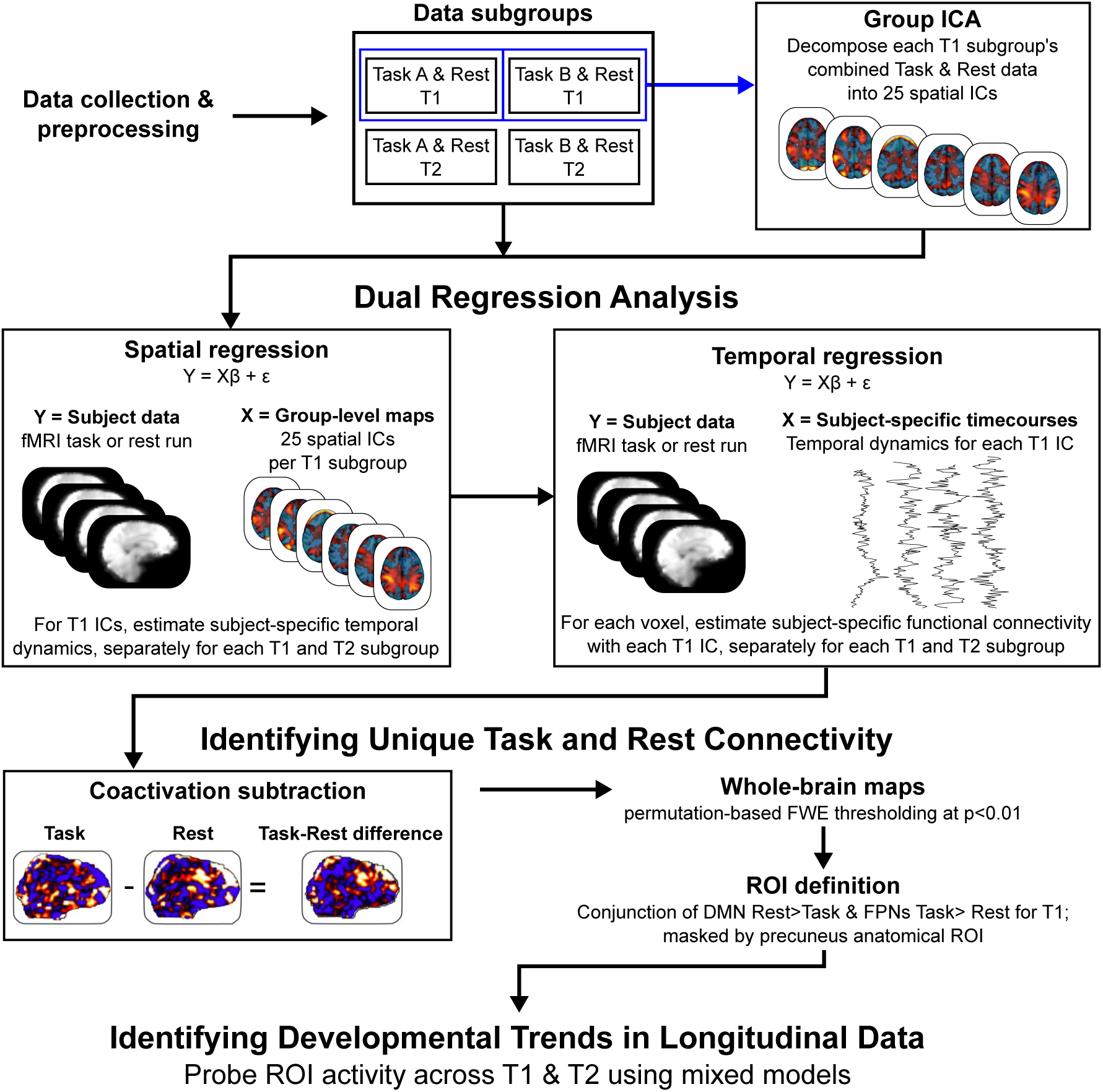
Schematic diagram of analytic approach (adapted from Utevsky, Smith, & Huettel, 2014). After data were collected and preprocessed, each task dataset was paired with the corresponding rest dataset for the same subgroup of participants. Data from two tasks across two developmental timepoints were used, resulting in four subgroups. Each T1 subgroup of combined task and rest data were entered into a group Independent Component Analysis (ICA), resulting in 25 spatial network maps for the T1 Task A + Rest data, and another 25 spatial network maps for the T1 Task B + Rest dataset. The network maps for each T1 Task subgroup were then entered in separate dual regression analyses for each corresponding T1 and T2 Task subgroup (e.g. T1 Task A + Rest network maps were used for a T1 Task A + Rest dual regression and a separate T2 Task A + Rest dual regression). This allowed us to quantify, for each participant, each voxel’s connectivity with each network while controlling for the other 24 networks. Each participant’s resting-state connectivity map was then subtracted from their task-state connectivity map, allowing us to examine within-participant connectivity differences for each of our networks of interest (default-mode network [DMN], left frontoparietal network [LFPN], and right frontoparietal network [RFPN]). The resulting task-rest difference maps were submitted to permutation-based thresholding to examine statistically significant differences. Finally, precuneus region of interest (ROI) analyses were conducted by submitting Task-Rest connectivity differences at both timepoints to mixed models including age-related parameters.

## Materials and Methods

### Participants and experimental tasks

Data for the current study were drawn from the first two timepoints of a large longitudinal imaging study (BrainTime) that was conducted at Leiden University in the Netherlands. At the first longitudinal timepoint (T1), data were collected from 299 participants (mean age: 14.15 years; age range: 8.01-25.95 years; 156 female). Data at the second longitudinal timepoint (T2) were collected approximately two years later from 254 of the original participants (mean age: 16.07 years; age range: 9.92-26.62; 131 female). While previous papers have published findings from the BrainTime study’s resting state data (Peters, Jolles, van Duijvenvoorde, Crone, & Peper, 2015; Peters, Peper, van Duijvenvoorde, Braams, & Crone, 2017; van Duijvenvoorde, Achterberg, Braams, Peters, & Crone, 2015; van Duijvenvoorde, Westhoff, de Vos, Wierenga, & Crone, submitted manuscript); and data from its two tasks separately (Braams, Güroǧlu, et al., 2014; Braams & Crone, 2016b, 2016a; Braams, Peper, Van Der Heide, Peters, & Crone, 2016; Braams, van Duijvenvoorde, Peper, & Crone, 2015; Braams, Peters, Peper, Güroğlu, & Crone, 2014; Peters & Crone, 2017; Peters, Braams, Raijmakers, Koolschijn, & Crone, 2014; Peters, Koolschijn, Crone, van Duijvenvoorde, & Raijmakers, 2014; Peters, van der Meulen, Zanolie, & Crone, 2017; Peters, Van Duijvenvoorde, Koolschijn, & Crone, 2016; Schreuders et al., 2018), this is the first study integrating all of the functional task and rest data from BrainTime’s first two longitudinal timepoints.

Written informed consent was obtained from adult participants while parent consent and participant assent was obtained from minor participants under a protocol approved by the institutional review board of Leiden University Medical Centre. Participants were screened for MRI contra-indications. All participants were right-handed, had no history of neurological or psychiatric disorders, and had their anatomical scans reviewed and cleared for incidental findings by a radiologist.

At both longitudinal timepoints, participants first completed a 5.1-minute resting-state scan in which they were instructed to lie still with their eyes closed but remain awake. Then, participants completed two runs of a feedback learning task, in which they learned associations between stimuli through positive and negative feedback (Task A; Peters, Braams, et al., 2014; Peters, Koolschijn, et al., 2014; Peters, van der Meulen, et al., 2017; Peters et al., 2016), followed by two runs of a self/other reward processing task, in which they guessed coin flip outcomes to win money for themselves or another person (Task B; Braams, Güroǧlu, et al., 2014; Braams & Crone, 2016a, 2016b, Braams et al., 2016, 2015; Braams, Peters, et al., 2014). Finally, anatomical images were obtained.

Data were split into four subgroups based on longitudinal timepoint and task (e.g. the original T1 TaskA+Rest subgroup consisted of all participants who, at T1, completed at least one run of Task A and the resting state run). Within each subgroup, data were excluded based on data quality concerns (see *Preprocessing below*) or for task performance indicating poor task engagement (performance less than three times the interquartile range in the feedback learning task; failing to make a response on >30% of trials in the reward processing task). These exclusion criteria left a final sample of 225 participants in the T1 Task A+rest subgroup, 197 participants in the T1 Task B+rest subgroup, 198 participants in the T2 Task A+rest subgroup, and 187 participants in the T2 Task B + rest subgroup. See Table 1 for demographics of each subgroup.

**Table 1:** Participant demographics for each subgroup.

| Group | Final N<br>(N female) | Mean age<br>(years) | Age range<br>(years) |
| --- | --- | --- | --- |
| T1 Task A + rest | 225 (116) | 14.58 | 8.01-25.95 |
| T1 Task B + rest | 197 (109) | 14.82 | 8.01-25.95 |
| T2 Task A + rest | 198 (102) | 16.58 | 10.02-26.62 |
| T2 Task B + rest | 187 (96) | 16.75 | 10.02-26.62 |

### Image acquisition

Scanning was performed on a 3-T Phillips Achieva MRI system using a standard whole-head coil. Functional scans were acquired using a T2*-weighted echo-planar imaging (EPI) sequence (TR = 2.2 s, TE = 30 ms, descending sequential acquisition of 38 axial slices, flip angle = 80°, FOV = 220 × 114.7 × 220 mm^3^, voxel size = 2.75 mm^3^). The resting state scan consisted of 142 volumes, each run of Task A consisted of 128 to 222 volumes, and each run of Task B consisted of 212 volumes. All functional runs included two initial dummy volumes to allow for signal equilibration. Experimental images were back-projected onto a screen that was viewed through a mirror.

Following functional scanning, a T1-weighted anatomical scan (TR = 9.76 ms, TE = 4.59 ms, flip angle = 8°, FOV = 224 × 168 × 177.3, 140 slices, voxel size = 1.17 × 1.17 × 1.2 mm, inversion time = 1050 ms) and a high-resolution EPI scan (TR = 2.2 s, TE = 30 ms, 84 slices, flip angle = 80°, FOV = 220 × 168 × 220 mm^3^, voxel size = 1.96 × 2 × 1.96 mm), were obtained to facilitate coregistration and normalization of functional data.

### Preprocessing

Data were preprocessed using FSL version 5.0.4’s (Smith et al., 2004; Woolrich et al., 2009) FMRIB’s Expert Analysis Tool (FEAT), including motion correction by realignment to the middle volume of each time series (Jenkinson & Smith, 2001), slice-time correction, removal of non-brain tissue (Smith, 2002), spatial smoothing using a Gaussian kernel of 6 mm full width at half maximum, grand mean intensity normalization, and high-pass temporal filtering (Gaussian-weighted least-squares straight line fitting with a 150s cut-off). Functional scans were first coregistered to the high-resolution EPI images, which were in turn registered to the T1 images, which were finally registered to the Montreal Neurological Institute (MNI) avg152 T1-weighted template using FSL’s Nonlinear Image Registration Tool (FNIRT).

For quality control, we examined 5 partially correlated measures of quality assurance for each run: 1) average signal-to-fluctuation-noise ratio (SFNR; Friedman and Glover, 2006), 2) average volume-to-volume motion, 3) maximum absolute motion, 4) percentage of volumes with framewise displacement greater than 0.5 mm (Power, Barnes, Snyder, Schlaggar, & Petersen, 2012), 5) percentage of outlier volumes with root-mean-square intensity difference relative to the reference volume (refRMS) greater than the 75^th^ percentile plus the value of 150% of the interquartile range of refRMS for all volumes in the run (i.e. standard boxplot threshold for outlier detection). Within each participant’s data for each task, we identified the “best” of the two runs as the run with the lowest percentage of refRMS outlier volumes and included only that run in subsequent analyses. Finally, within each task’s best runs and within the resting state data, we excluded the 95^th^ percentile of worst runs for each data quality metric and/or >= 3mm of motion and/or >= 10% poor quality volumes, whichever criteria was strictest (see *Participants and experimental tasks* section for final included sample sizes). This left us with one task and one rest run per participant in each analysis subgroup (i.e. each participant in the T1 Task A+rest subgroup contributed one T1 Task A run and their T1 resting state run to the analyses). Because each analysis subgroup drew from the same initial participant pool but faced slightly different percentile-based exclusion criteria, the exact makeup of each analysis subgroup included overlapping but non-identical participants.

As even mild motion artifacts can distort connectivity analyses (Power et al., 2012; Satterthwaite et al., 2012), we implemented additional analyses to correct for motion issues. We regressed out variance tied to six motion parameters (rotations and translations along the three principal axes). Furthermore, for every run, we regressed out all volumes with framewise displacement greater than 0.5 mm and all refRMS outlier volumes. Though this is not identical to the scrubbing procedure of Power and colleagues (2012), it accomplishes the same goal of removing signal discontinuities and spurious effects of head motion that cannot be accounted for by conventional motion regression.

### Independent Component Analysis

T1 data for each task+rest subgroup were run through separate probabilistic group independent component analysis (ICA; Beckmann & Smith, 2004). Each ICA had two inputs per subgroup participant: one resting-state scan and one task-based scan. Input data were first downsampled to 3mm isotropic resolution using a 12-parameter affine transformation implemented in FSL’s Linear Image Registration Tool (FLIRT; Jenkinson & Smith, 2001) to reduce data-processing demands. Through FSL’s Multivariate Exploratory Linear Optimized Decomposition into Independent Components (MELODIC), data were voxel-wise demeaned and normalized, whitened, and projected into a 25-dimensional subspace (the number of selected components was based on Utevsky et al., 2014), resulting in 25 independent components (ICs) per subsample.

From each T1 task+rest subgroup’s ICA output, we identified the ICs corresponding to our three networks of interest (DMN, RFPN, and LFPN) as those with the highest spatial correlation to the canonical network maps of Smith and colleagues (2009) (Figure 2A).

**Figure 2.**
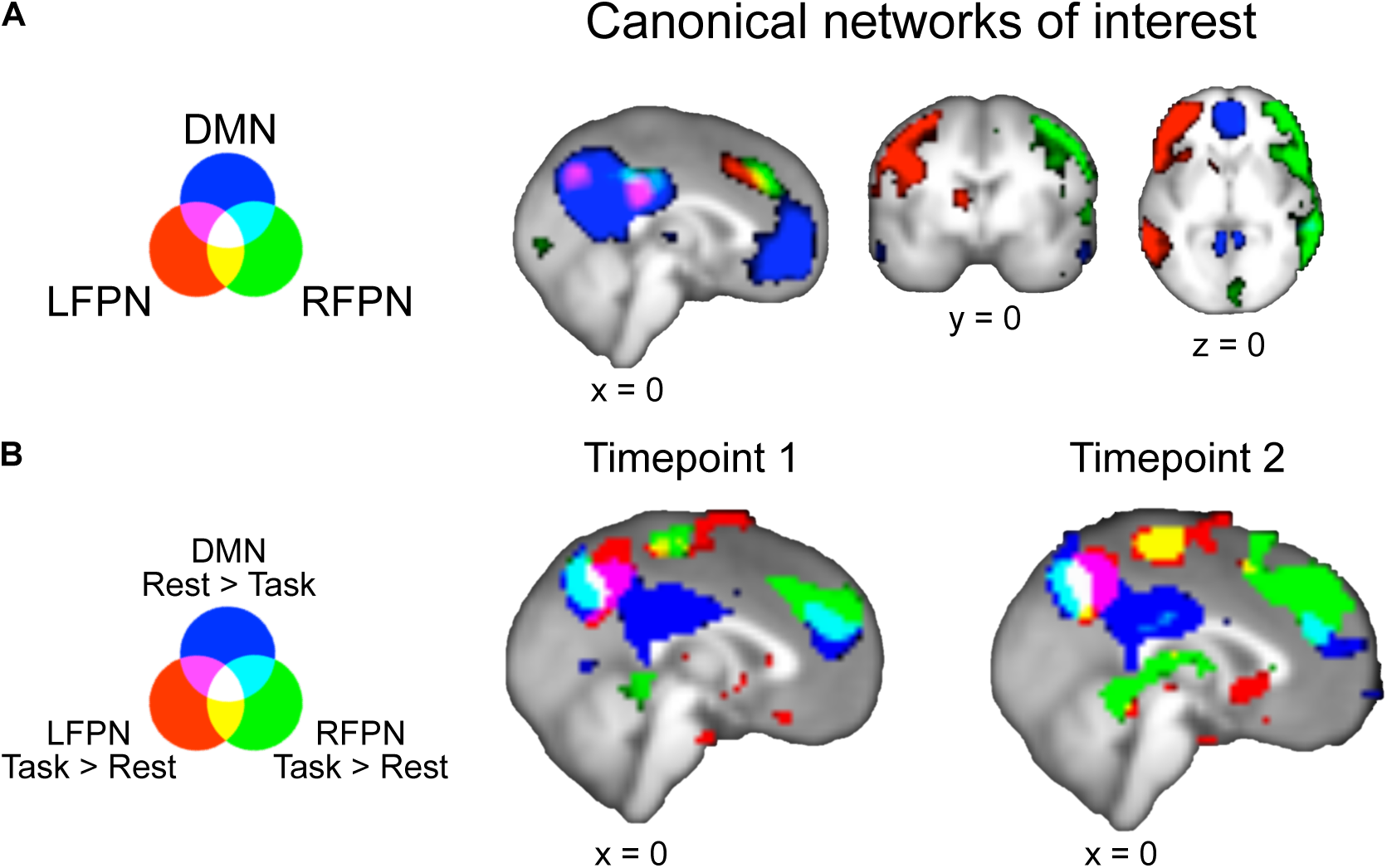
Task versus rest connectivity with default mode network (DMN) and left and right frontoparietal networks (LFPN and RFPN). **A.** Canonical networks of interest from Smith et al., 2009, downsampled to 3mm resolution. Images thresholded at 2.3 < *z* < 4. **B.** Task vs Rest connectivity for the three canonical networks of interest. rest>task connectivity with DMN and task>rest connectivity with LFPN and RFPN show a conjunction over precuneus (white). FWE-corrected *p*<0.01 with threshold-free cluster enhancement

### Dual-Regression Analysis

We examined changes in connectivity between task and rest states by submitting the network maps of each group’s task and rest runs to dual-regression analyses (as in Utevsky et al., 2014). Dual-regression analysis quantifies voxelwise connectivity for each IC while controlling for the other ICs (Filippini et al., 2009; Nickerson et al., 2017). Each dual-regression analysis comprised two stages (Figure 1). First, each IC map was regressed onto each run’s functional dataset, resulting in run-specific time courses for each IC. Second, those resulting timecourses were then regressed onto each run’s functional data to estimate each voxel’s connectivity with each IC while controlling for the other 24 ICs.

Each task+rest group at T1 used its own ICs for its own separate dual-regression analysis. So that the ICs entering the dual regression were consistent across longitudinal timepoints, T1 ICs were used for their corresponding T2 subgroups’ dual-regression. In other words, T1’s TaskA+Rest ICs were used for the dual-regression analysis of T1 TaskA+Rest and the dual-regression analysis for T2 TaskA+Rest, while T1’s TaskB+Rest ICs were used for the dual-regression of T1 TaskB+Rest and the dual-regression analysis for T2 TaskB+Rest.

### Task-Rest General Linear Model

To investigate differences in task and rest connectivity within each participant, each participant’s resting-state connectivity map was subtracted from their task-state connectivity map separately for each task (Task A > Rest; Task B > Rest), for each of the three networks of interest (DMN, LFPN, and RFPN) and at each longitudinal timepoint (T1 and T2). The resulting difference maps indicating task-minus-rest changes in connectivity with each network were entered into separate group-level general linear models for each task+rest subgroup, network, and longitudinal timepoint. To further control for spurious motion-related results that could arise from participants moving more during task than rest—above and beyond motion controls implemented during preprocessing—each model contained additional subject-level nuisance regressors representing individual differences in motion between task and rest: 1) difference in average signal-to-fluctuation-noise ratio, 2) difference in average volume-to-volume motion, 3) difference in percentage of volumes with framewise displacement greater than 0.5 mm, and 4) difference in percentage of refRMS outlier volumes. Finally, we included 5) an additional nuisance regressor to account for gender (Filippi et al., 2013; Smith et al., 2014). All nuisance regressors were demeaned.

We generated bidirectional contrasts comparing task and rest states in each of the three networks of interest for each task+rest subgroup at each longitudinal timepoint. Statistical significance was determined using Monte Carlo permutation-based statistical thresholding with 10,000 permutations, family-wise error corrected for multiple comparisons across the whole brain (Winkler, Ridgway, Webster, Smith, & Nichols, 2014). Activation clusters were estimated using threshold-free cluster enhancement (Smith & Nichols, 2009).

We conducted conjunction analyses across both tasks, separately for each longitudinal timepoint (i.e. conjunction of T1 TaskA>rest with T1 TaskB>rest; conjunction of T2 TaskA>rest with T2 TaskB>rest) for each network of interest using the minimum statistic (Nichols et al. 2005). This allowed us to examine neural connectivity during task states in general, rather than connectivity specific to Task A or Task B. These conjunction analyses resulted in a task>rest and a rest>task connectivity map for each network of interest (DMN, LFPN, and RFPN) and at each longitudinal timepoint.

### Region of Interest (ROI) Identification

In order to restrict subsequent age-related analyses to our *a priori* region of interest, we identified a precuneus ROI by masking the T1 conjunction of task>rest connectivity with LFPN, task>rest connectivity with RFPN, and rest>task connectivity with DMN (see Figure 2B) with the Harvard-Oxford atlas’s anatomical precuneus ROI thresholded at 70% (Desikan et al., 2006). Participants’ task and rest connectivity parameters were then extracted from this precuneus ROI for each longitudinal timepoint and network.

### Experimental Design and Statistical Analysis

Age-related differences in connectivity parameters were further probed across data from both longitudinal timepoints using a mixed model approach in R with the package *nlme* (Pinheiro, Bates, DebRoy, Sarkar, & Team, 2018). Mixed models (also known as hierarchical linear models, multilevel models, or random-effects models) as applied to longitudinal datasets allow longitudinal timepoints to be nested within participants by modeling participant identity as a random effect. ROI mixed models were run on connectivity parameters combined across T1 and T2, separately for each Task (e.g. T1 & T2 Task A + Rest data in one set of analyses; T1 & T2 Task B + Rest data in a separate set of analyses). This allowed us to self-replicate any developmental changes in task-versus-rest connectivity across two different tasks.

To test for significant developmental differences, we first fit a null, intercept-only model including a fixed and random intercept. We then compared the intercept-only model to three different age-related models: one with a mean-centered linear continuous age term to test for monotonic age-related changes, one with mean-centered linear and quadratic age terms to test for additional quadratic age-related changes (e.g. developmental peaks or troughs), and one with mean-centered linear, quadratic, and cubic age terms to test for additional cubic age-related changes (e.g. developmental changes that emerge and then stabilize; Madhyastha et al., 2017). Likelihood ratio tests between the intercept-only, linear, quadratic, and cubic models were used to determine whether the age-related models with significant age parameters significantly improved model fit over the intercept-only model or the next simplest model with a significant age parameter.

In order to rule out overall task performance or engagement as a confound for any significant age-related findings, we ran additional models adding a metric of task engagement as an additional regressor to any model with significant age-related findings that may have been driven by neural activity during task. The metric for Task A was participants’ learning rates, as measured by the percentage of trials in which feedback was successfully used on the subsequent trial (Peters, Braams, et al., 2014; Peters et al., 2016). The metric for Task B was participants’ self-reported liking of winning money for self, reported at the end of the scanning session (Braams, Güroǧlu, et al., 2014; Braams, Peters, et al., 2014; Braams et al., 2015). Task B self-report metrics were not collected from 87 participants at T1 and 6 participants at T2, so task engagement control analyses were run with 353 datapoints from 248 participants for Task B (see Table 2).

**Table 2:** Number of participants and datapoints in mixed model analyses.

| T1 & T2 Task + Rest subgroup | N participants | N datapoints |
| --- | --- | --- |
| Task A + Rest:<br>All participants | 265 | 423 |
| Task B + Rest:<br>All participants | 253 | 384 |
| Task A + Rest:<br>Task engagement analyses | 265 | 423 |
| Task B + Rest:<br>Task engagement analyses | 248 | 353 |

Because self-reported task engagement metrics were not collected from a subset of Task B participants, task engagement control analyses for Task B were run with fewer participants and datapoints than the age-only models.

## Results

### Precuneus connectivity distinguishes task and rest states across development

The whole-brain conjunction analyses across both tasks at each longitudinal timepoint show that precuneus is both significantly more connected with both LFPN and RFPN at task than at rest, and significantly more connected with DMN at rest than during task (Figure 2B, Table 3). Thus, in our cross-sectional samples at both T1 and T2, we replicate Utevsky and colleagues’ (2014) finding that precuneus connectivity distinguishes between tasks and rest states in adults via varying connectivity with DMN (rest>task) and LFPN (task>rest). Notably, we also extend this finding to connectivity with RFPN (task>rest), and to cross-sectional neural data from participants between the ages of 8 and 26.

**Table 3:** Regions exhibiting task- and rest-dependent connectivity changes with DMN and FPNs.

| Longitudinal<br>timepoint | Probabilistic<br>anatomical label | X | Y | Z | *p* value | cluster<br>extent |
| --- | --- | --- | --- | --- | --- | --- |
| T1 | precuneus cortex<br>(86%) | 3 | -63 | 33 | <0.001 | 152 |
|  | inferior lateral<br>occipital cortex<br>(61%), occipital<br>fusiform gyrus<br>(9%) | -45 | -75 | -9 | <0.001 | 108 |
|  | postcentral gyrus<br>(51%), precentral<br>gyrus (30%) | -60 | -6 | 24 | <0.001 | 101 |
|  | insular cortex (74%),<br>frontal orbital<br>cortex (8%) | -36 | 15 | -12 | 0.002 | 12 |
| T2 | precuneus cortex<br>(26%), cuneal<br>cortex (18%),<br>supracalcarine<br>cortex (17%),<br>intracalcarine<br>cortex (9%) | 12 | -66 | 21 | <0.001 | 179 |
|  | inferior lateral<br>occipital cortex<br>(50%), superior<br>lateral occipital<br>cortex (11%) | -39 | -81 | 6 | <0.001 | 75 |
|  | superior lateral<br>occipital cortex<br>(67%), precuneus<br>(1%) | 30 | -69 | 51 | <0.001 | 71 |
|  | precentral gyrus<br>(23%), postcentral<br>gyrus (19%),<br>central opercular<br>cortex (4%) | -57 | -6 | 18 | 0.003 | 63 |
|  | frontal pole (87%) | -24 | 60 | 0 | <0.001 | 35 |
|  | inferior lateral<br>occipital cortex<br>(39%), occipital<br>fusiform gyrus<br>(19%) | -42 | -72 | -12 | <0.001 | 26 |

Regions at the conjunction of left frontoparietal network (LFPN) task > rest, right frontoparietal network (RFPN) task > rest, and default mode network (DMN) rest > task, FWE-corrected $p < 0.01$ (see *Methods*) with a cluster extent of at least 10 voxels. Probabilistic anatomical labels refer to the likelihood that the listed max voxel coordinates are within the given Harvard-Oxford Cortical Structural Atlas region.

The whole brain conjunction also indicated that lateral occipital cortex and pre-/postcentral gyrus exhibited a similar connectivity profile to precuneus, though we note that precuneus was the largest conjunction cluster at both longitudinal timepoints (Table 3). As we *a priori* hypothesized that precuneus would exhibit state-dependent connectivity changes (Utevsky et al., 2014), subsequent ROI analyses to probe developmental connectivity changes were conducted within our conjunction result, masked by a precuneus anatomical ROI (see *Materials and Methods: ROI Identification*).

### Task/Rest connectivity differences between precuneus and LFPN increase with age

We applied mixed models to the longitudinal precuneus connectivity parameters extracted from our precuneus ROI. This allowed us to test for age-related changes in task-rest connectivity between the precuneus and each of the three networks of interest, while accounting for the repeated measures in our longitudinal data. Separate mixed models were applied to TaskA+Rest data and to TaskB+Rest data, which allowed us to use two different tasks to 1) self-replicate any developmental findings across tasks and 2) show that such findings are generalizable to task-state rather than specific to a particular task.

We found a significant linear effect of age in task>rest precuneus connectivity with LFPN in both of the tasks (β_LinearAge_ = 0.56, *p* = 0.0016 for Task A subgroup; (β_LinearAge_ = 0.55, *p* < 0.001 for Task B subgroup), such that increasing age was associated with greater task-minus-rest differences in precuneus-LFPN connectivity (Figure 3A). For both tasks, adding a linear age term significantly improved model fit over the intercept-only model (χ^2^(1) = 10.19, *p* = 0.0014 for Task A subgroup; χ^2^(1) = 12.65, *p* < 0.001 for Task B subgroup). Additional quadratic and cubic age regressors were not significant in models for both tasks (all *p*s > 0.05).

**Figure 3:**
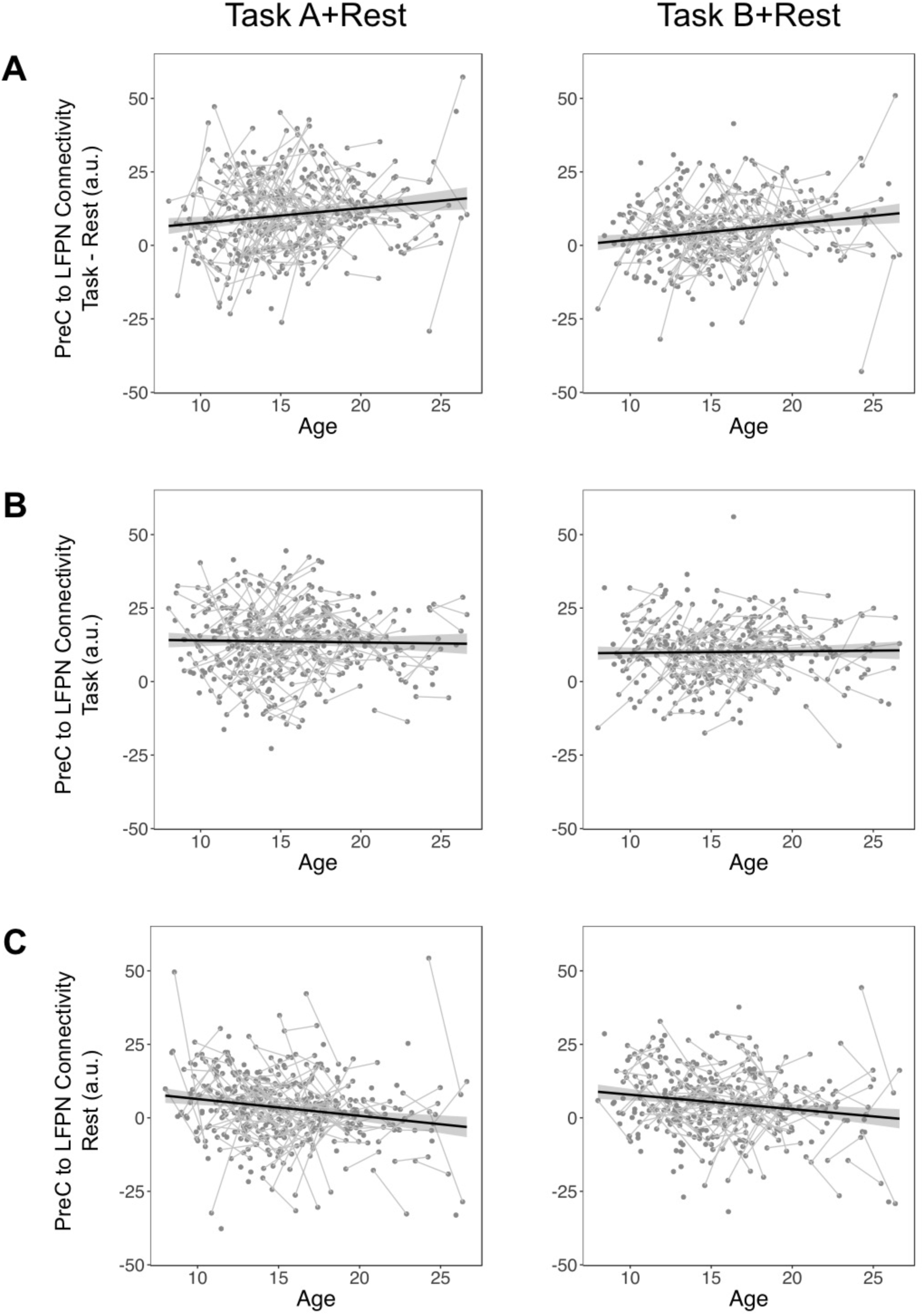
Precuneus-left frontoparietal network (LFPN) connectivity changes across development at rest but not task. **A.** Across both tasks, task-rest connectivity differences between precuneus and LFPN significantly linearly increased with age. Further probing task-connectivity and rest-connectivity revealed **B.** no significant age-related relationships between precuneus-LFPN connectivity at task but **C.** a significant linear age-related decrease in precuneus-LFPN connectivity at rest. Note that the resting state data in both C panels were drawn from the full set of participants. Each C panel reflects overlapping but different samples that were submitted to separate dual regression analyses. Connected lines link longitudinal datapoints from the same participant. Shaded areas represent 95% confidence intervals around linear best-fit lines.

We next ran control analyses to examine whether these effects could be attributed to overall task performance or engagement. When learning rate, as a metric of task performance/engagement, was added to the linear age model for Task A (feedback learning task), age remained a significant predictor (β_LinearAge_ = 0.51, *p* = 0.0070), but learning rate was not a significant predictor (β_LearningRate_ = 0.12, *p* = 0.39) of task>rest precuneus-LFPN connectivity. When self-reported reward liking, as a metric of task performance/engagement, was added to the linear age model for Task B (reward processing task), age remained a significant predictor (β_LinearAge_ = 0.52, *p* = 0.0037) but reward liking was not a significant predictor (β_RewardLiking_ = - 0.26, *p* = 0.41) of task>rest precuneus-LFPN connectivity. Thus, our age-related changes in connectivity cannot be attributed to age-related changes in overall task performance or engagement.

We found no significant age-related changes in task-minus-rest precuneus connectivity with DMN and RFPN that replicated across both tasks. There was a significant cubic effect of age on task-minus-rest connectivity between precuneus and DMN in the Task A subgroup (β_CubicAge_ = 0.20, *p* = 0.035), but the model with linear, quadratic, and cubic age regressors did not fit significantly better than the intercept only model (χ^2^(3) = 4.54, *p* = 0.21). Furthermore, this cubic effect of age on precuneus-DMN connectivity did not replicate in the Task B subgroup (β_CubicAge_ = - 0.0027, *p* = 0.80). All other linear, quadratic, and cubic age effects were non-significant for task-minus-rest connectivity between precuneus and DMN, and precuneus and RFPN (all *p*s > 0.05). Thus, although precuneus was significantly more connected to DMN for rest>task, and significantly more connected to RFPN for task>rest across the sample and at both timepoints, no developmental changes were observed in this connectivity.

### Connectivity between precuneus and LFPN diminishes with age during rest but not task

To further probe whether the age-related linear increase in task-rest precuneus connectivity with LFPN was due to increased connectivity during task or decreased connectivity during rest, we ran additional mixed models on the connectivity parameters between precuneus and LFPN at task and at rest. For both Task A and Task B, there was no significant linear effect of age upon precuneus-LFPN connectivity during task (β_LinearAge_ = 0.80, *p* = 0.63 for Task A; β_LinearAge_ = 0.088, *p* = 0.55 for Task B; Figure 3B). Additional models with task engagement and linear age as predictors of precuneus-LFPN connectivity during task found age and task engagement to be non-significant predictors in both tasks, and task engagement alone also failed to significantly predict precuneus-LFPN connectivity during task (all predictor *p*s > 0.05). Thus, precuneus connectivity to the LFPN task-based network neither varied by age nor by task nor by task engagement.

There was, however, a significant linear effect of age during rest (β_LinearAge_ = - 0.57, *p* < 0.001 for Task A subgroup; β_LinearAge_ = −0.50, *p* = 0.0020 for Task B subgroup), such that increasing age was associated with reduced connectivity between precuneus and LFPN at rest (Figure 3C; we reiterate here that the Task A and Task B subgroups represent overlapping but non-identical samples drawn from the full participant set and submitted to separate dual regression analyses). For both subgroups, adding a linear age term significantly improved model fit over the intercept-only model (χ^2^(1) = 12.12, *p* < 0.001 for Task A subgroup; χ^2^(1) = 9.86, *p* = 0.0017 for Task B subgroup). Additional quadratic age regressors were non-significant in the models with linear and quadratic age (β_QuadraticAge_ = 0.014, *p* = 0.67 for Task A subgroup; β_QuadraticAge_ = −0.0086, *p* = 0.78 for Task B subgroup).

In the Task A subgroup, the model with linear, quadratic, and cubic age regressors resulted in a non-significant effect of linear age (β_LinearAge_ = −0.030, *p* = 0.91) and significant effects of quadratic (β_QuadraticAge_ = 0.085, *p* = 0.045) and cubic (β_CubicAge_ = −0.017, *p* = 0.0089) age. This cubic age model fit significantly better than the linear age model for the Task A subgroup (χ^2^(2) = 7.16, *p* = 0.028). In the Task B subgroup, however, the cubic age model resulted in non-significant regressors for linear, quadratic, and cubic age (β_LinearAge_ = −0.11, *p* = 0.68, β_QuadraticAge_ = 0.034, *p* = 0.40, and β_CubicAge_ = −0.011, *p* = 0.087, respectively), and the cubic age model did not fit significantly better than the linear age model (χ^2^(2) =3.07, *p* = 0.22). These results suggest that linear age may be a more consistently parsimonious predictor of developmental changes in precuneus-LFPN connectivity at rest.

## Discussion

In this study, we investigated the state-dependent functional connectivity of the precuneus across development in a large cross-sectional and longitudinal sample of participants between the ages of 8 and 26. Previous work has shown that the precuneus serves as a unique hub distinguishing between task and rest states in the adult brain (Utevsky et al., 2014), yet little has been known about the role of precuneus in the developing brain. Using dual regression analyses to track network-based functional connectivity across the sample, we show that the precuneus exhibits both greater functional connectivity with two task-based networks (RFPN and LFPN) during task compared to rest, as well as greater functional connectivity with the rest-based DMN during rest compared to task. This result replicated across two different tasks and at two longitudinal timepoints. Thus, the role of precuneus in mediating between task and rest states is evident throughout development from childhood through early adulthood.

We next examined whether the mediating function of the precuneus strengthens or matures over development by searching for age-related changes in state-dependent functional connectivity between precuneus and LFPN, RFPN, and DMN. Across two distinct tasks, we found a significant linear age-related increase in state-dependent functional connectivity between precuneus and LFPN, such that task>rest connectivity between precuneus and LFPN significantly linearly increased with age from childhood to adulthood. We determined that the relationship between age and task>rest precuneus-LFPN connectivity was driven by an age-related decrease in precuneus-LFPN connectivity during rest. Such age-related increasing segregation between LFPN and a region of DMN during rest is consistent with prior work showing age-related increases in network segregation in both structural (Baum et al., 2017) and resting state data (Fair et al., 2007, 2008; Sherman et al., 2014; Uddin et al., 2011). We note that our developmental findings are strengthened by BrainTime’s accelerated longitudinal design, in which both cross-sectional and longitudinal effects can be captured.

In contrast to the observed developmental connectivity changes during rest, task-related connectivity between precuneus and LFPN was continuously high across ages and showed no developmental changes. This suggests that age-related increases in network segregation shown during rest (Fair et al., 2007, 2008; Sherman et al., 2014; Uddin et al., 2011) may not occur during task and/or do not apply to precuneus-LFPN connectivity. Finally, we found that precuneus’s state-dependent connectivity with DMN (rest>task) and RFPN (task>rest) remained developmentally stable, with no significant effect of linear, quadratic, and/or cubic age.

Notably, we did not find task performance or engagement to significantly predict task>rest precuneus-LFPN connectivity after accounting for the effect of age. Furthermore, we found no significant relationship between individuals’ task performance or engagement and their precuneus-LFPN connectivity during task. This is surprising, given that prior work has shown that different tasks differentially affect segregation and integration between neural networks (Cohen & D’Esposito, 2016; Khambhati, Medaglia, Karuza, Thompson-Schill, & Bassett, 2018), and that task performance affects the FPN’s overall activation and functional connectivity throughout the brain (Cole et al., 2013; Dwyer et al., 2014; Satterthwaite et al., 2013). Thus, our findings highlight the unique role of the precuneus, which tracks whether the brain is engaged in a task state generally, with no observed differences between the two tasks in the BrainTime dataset. Future work could investigate whether precuneus-LFPN connectivity is moderated by task engagement for more variably engaging/demanding tasks (e.g. N-back tasks with varying N as in Cohen & D’Esposito, 2016; Satterthwaite et al., 2013). Application of network-based psychophysiological interaction approaches (Utevsky, Smith, Young, & Huettel, 2017) can also be used to determine whether the timecourse of precuneus-LFPN connectivity is modulated by moment-to-moment changes in task demands (e.g. precuneus-LFPN connectivity for high demand > low demand blocks within the same run).

We note that the present study replicates the previous findings of Utevsky and colleagues’ (2014) study of adult neural data, and extends them in three significant ways. First, we extend the findings to a large cross-sectional *and* longitudinal developmental sample, which allowed us to show that the role of precuneus as a functional core of the DMN is in place in childhood. The developmental trajectory of task>rest precuneus-LFPN connectivity suggests, however, that younger populations than those in our sample (e.g., children younger than 8 years) may exhibit no differences in precuneus-LFPN connectivity for task versus rest states. Thus, future work could examine precuneus connectivity in even younger participants in order to determine if this is, in fact, the case. Second, our work further extends the precuneus connectivity finding to two tasks different from those used in the adult study (Utevsky et al., 2014). This strengthens the hypothesis that precuneus mediates between rest and task generally, regardless of the specific task. Third, our study complements prior work: While the previous adult study collected the resting-state scan last, after the completion of the three tasks (Utevsky et al., 2014), our study collected the resting-state scan first, prior to the task runs. This avoids the concern that recent exposure to a task may alter subsequent resting-state connectivity (Stevens, Buckner, & Schacter, 2010; Tung et al., 2013; Waites, Stanislavsky, Abbott, & Jackson, 2005). Thus, the two studies together suggest that the precuneus’s unique role in mediating between task and rest states holds, regardless of task versus rest order.

Prior studies investigating developmental changes in whole-brain connectivity patterns have often used graph-theoretic approaches that examine connectivity strength between distributed nodes (Fair et al., 2007, 2008; see Ernst, Torrisi, Balderston, Grillon, & Hale, 2015; Power, Fair, Schlaggar, & Petersen, 2010; M. C. Stevens, 2016 for review). Such developmental work, however, generally investigates connectivity during rest rather than during task (but see Joseph et al., 2012) and do not directly compare across task and rest states, as in the present study. Given that resting state graph theory analyses have also implicated precuneus as a key functional hub (Tomasi & Volkow, 2010), combining dual-regression and task-versus-rest analyses with graph theory approaches could be complementary and yield further insight as to how neural hubs affect processing state.

Additional future studies could also use structural neuroimaging techniques, such as diffusion tensor imaging, to determine whether developmental changes in precuneus-LFPN functional connectivity are reflected in structural changes in the developing brain. Converging evidence from non-human primate anatomical tracing studies (Leichnetz, 2001) and resting-state functional connectivity work in humans and non-human primates (Margulies et al., 2009) suggest that there are cortical projections between precuneus and regions of the LFPN, including lateral prefrontal cortex and lateral parietal cortex (Cavanna & Trimble, 2006). Future work should examine whether and how such structural connections between precuneus and LFPN underlie state-dependent differences in functional connectivity, and how they may change across development.

It is important to understand developmental changes in task and rest state connectivity, as altered connectivity has been reported for numerous psychological disorders (see Cohen, 2017; Greicius, 2008 for review). For example, a maturational lag in DMN to FPN connectivity has been associated with attention-deficit/hyperactivity disorder (Sripada, Kessler, & Angstadt, 2014), and abnormal resting-state functional connectivity between DMN and FPN has been found in patients with obsessive compulsive disorder (Stern, Fitzgerald, Welsh, Abelson, & Taylor, 2012) and with impaired consciousness (Long et al., 2016). Future work could investigate whether precuneus’s task- and rest-based connectivity is altered in atypical development, and whether any such deviations from typical connectivity development correspond to behavioral or psychological impairments.

In this study, we demonstrate that precuneus plays a key role in mediating between task and rest states via connectivity with DMN and FPNs across typical development. These results underscore the unique nature of an enigmatic brain region that has been implicated in processes as varied as memory, self-processing, decision-making, and even consciousness (Cavanna & Trimble, 2006), while also pointing to future targets for understanding changes in neural connectivity in typical and atypical development.

## Acknowledgments

This work was supported by an innovative ideas grant of the European Research Council (ERC-2010-StG-263234) to E.A.C. and a Graduate Research Opportunities Worldwide Fellowship from the National Science Foundation and the Netherlands Organisation for Scientific Research to R.L.

